# Tissue pressure and cell traction compensate to drive robust aggregate spreading

**DOI:** 10.1101/2020.08.29.273334

**Authors:** M. S. Yousafzai, V. Yadav, S. Amiri, M.F. Staddon, A. P. Tabatabai, Y. Errami, G. Jaspard, S. Amiri, S. Banerjee, M. Murrell

## Abstract

In liquid droplets, the balance of interfacial energies and substrate elasticity determines the shape of the droplet and the dynamics of wetting. In living cells, interfacial energies are not constant, but adapt to the mechanics of their environment. As a result, the forces driving the dynamics of wetting for cells and tissues are unclear and may be context specific. In this work, using a combination of experimental measurements and modeling, we show the surface tension of cell aggregates, as models of active liquid droplets, depends upon the size of the aggregate and the magnitude of applied load, which alters the wetting dynamics. Upon wetting rigid substrates, traction stresses are elevated at the boundary, and tension drives forward motion. By contrast, upon wetting compliant substrates, traction forces are attenuated, yet wetting occurs at a comparable rate. In this case, capillary forces at the contact line are elevated and aggregate surface tension contributes to strong outward, pressure-driven cellular flows. Thus, cell aggregates adapt to the mechanics of their environments, using pressure and traction as compensatory mechanisms to drive robust wetting.

---

The balance between cell-cell and cell-extracellular matrix (ECM) forces determines the collective motion of cells, influencing essential life processes, including embryonic development ^1, 2, 3, 4^ and spreading of cancer ^5, 6, 7, 8^. The propensity for tissues to flow^9, 10 11^, fuse ^12^ and distribute their stresses at the surface^13^ has led to modeling tissue as a fluid. Thus, cell-cell and cell-ECM interactions are abstracted as interfacial energies in a liquid droplet ^14, 15, 16, 17, 18^. Thermodynamic models such as the Differential Adhesion Hypothesis (DAH) ^2, 12, 19^ and mechanical models including the Differential Interfacial Tension Hypothesis ^20, 21 22^ describe how the balance between interfacial energies or tensions impacts the equilibrium configurations of tissues. However, there is no sufficient framework to describe the dynamic motions of cells and tissues from interfacial tensions out-of-equilibrium.

Active stresses are generated within the cell cytoskeleton, due to the non-equilibrium activity of molecular motors that convert chemical energy into mechanical work. Motor activity is associated with non-Boltzmann binding statistics and network fluctuations^23, 24, 25^, and the accumulation of active stress leads to the amplification of system dynamics and mechanical non-linearities^26, 27^. As a result, interfacial tensions in cells and simple tissues may not be constant ^28^, but vary with applied mechanical load ^29^, boundary conditions^30^, and system size^27^. Thus, the balance of these tensions with substrate elasticity may yield driving forces for wetting that depend upon the mechanics of the ECM. However, to date the dependence of surface tension on motor activity or extrinsic factors, and the influence of an adaptive surface tension of the dynamics of wetting are unknown.

In this work, we use a combination of mechanical experiments and physical models to understand the impact of active stress on the balance of interfacial tensions and substrate elasticity, and how this balance influences the dynamics of wetting of cell aggregates. First, we quantify the differences in wetting dynamics subject to different substrate elasticities. Then, we explain these dynamics through comparisons of the energy scales between measured aggregate surface tensions and substrate deformations, as determined by elasto-capillarity. Based on the comparison of energy scales, we identify distinct mechanisms that drive wetting for different substrate elasticities. We then confirm these mechanisms by computational and theoretical modeling. These results establish novel modes of collective cell migration and suggest mechanical adaptation as a means of regulating motion.

## Aggregate spreading is tissue size and substrate rigidity dependent

Aggregates of S180 Sarcoma cells, approximately 100 μm in diameter, are added to fibronectin-coated polyacrylamide gels that vary in rigidity (E) with constant surface adhesion ([fibronectin] = 1 mg/mL) (Fig 1A, Supplementary Figs 1, 2, Supplementary Tables 1, 2, 3). Aggregates, with initial radius R_0_ and maximal cross-sectional area A_0_, are visualized using Differential Interference Contrast (DIC) as well as fluorescent F-actin (Ftractin-EGFP) microscopy. Aggregates adhere and spread homogenously (Supplementary Fig 3, Supplementary Video 1, 2), expanding a monolayer, resembling the wetting of a liquid droplet and the expansion of a precursor film (Fig 1B) ^31 32, 33^. The aggregate of maximal cross-sectional area A_0_ exhibits ortho-radial F-actin organization and deformation of cell nuclei which varies with distance from the aggregate center (Fig 1C-E). The monolayer however, displays F-actin organization and focal adhesion maturation, quantified by the nematic order parameter 〈*q*〉 and focal adhesion size A_adhesion_ respectively, which depends upon substrate stiffness (Fig 1F-M, Methods). Focal adhesions are small and numerous on soft substrates (E =0.7 kPa) but are large and less numerous on rigid substrates (E = 40 kPa, glass). Consistent with alterations in cytoskeletal organization and adhesions, the spreading dynamics of the monolayer indicates subtle substrate stiffness dependence (Fig 1N-P). Measuring the monolayer strain (ε(t) = A(t)/A_0_) over time (t) shows a linear expansion of the monolayer on rigid substrates and a weakly exponential expansion on soft substrates (Fig 1O, Supplementary Fig 4). However, both the strain ε and strain rate 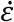 (defined as 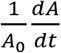) are similar in magnitude for A>A_0_ (Fig 1O, inset). Comparing the goodness of fit for both linear and exponential approximations (R^2^), the change from linear to exponential spreading dynamics occurs at approximately 10 kPa (Fig 1P). Finally, we show that on soft substrates (E = 0.7 kPa) strain rate 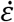 depends inversely on the size of aggregate (Fig 1Q), but no such relationship is observed on hard substrates (E = 40 kPa), or glass where the fit is non-significant. Douezan et al. have shown that balancing the rates of energy gained by adhesion and energy lost by slippage gives rise to a spreading behavior where the adhesion area increases linearly with time ^34^. However, a nonlinear behavior may suggest different mechanisms during motion. To further inform mechanisms of spreading we measure the spatiotemporal patterns of velocity and stress exerted on the substrate.

**Figure 1:**
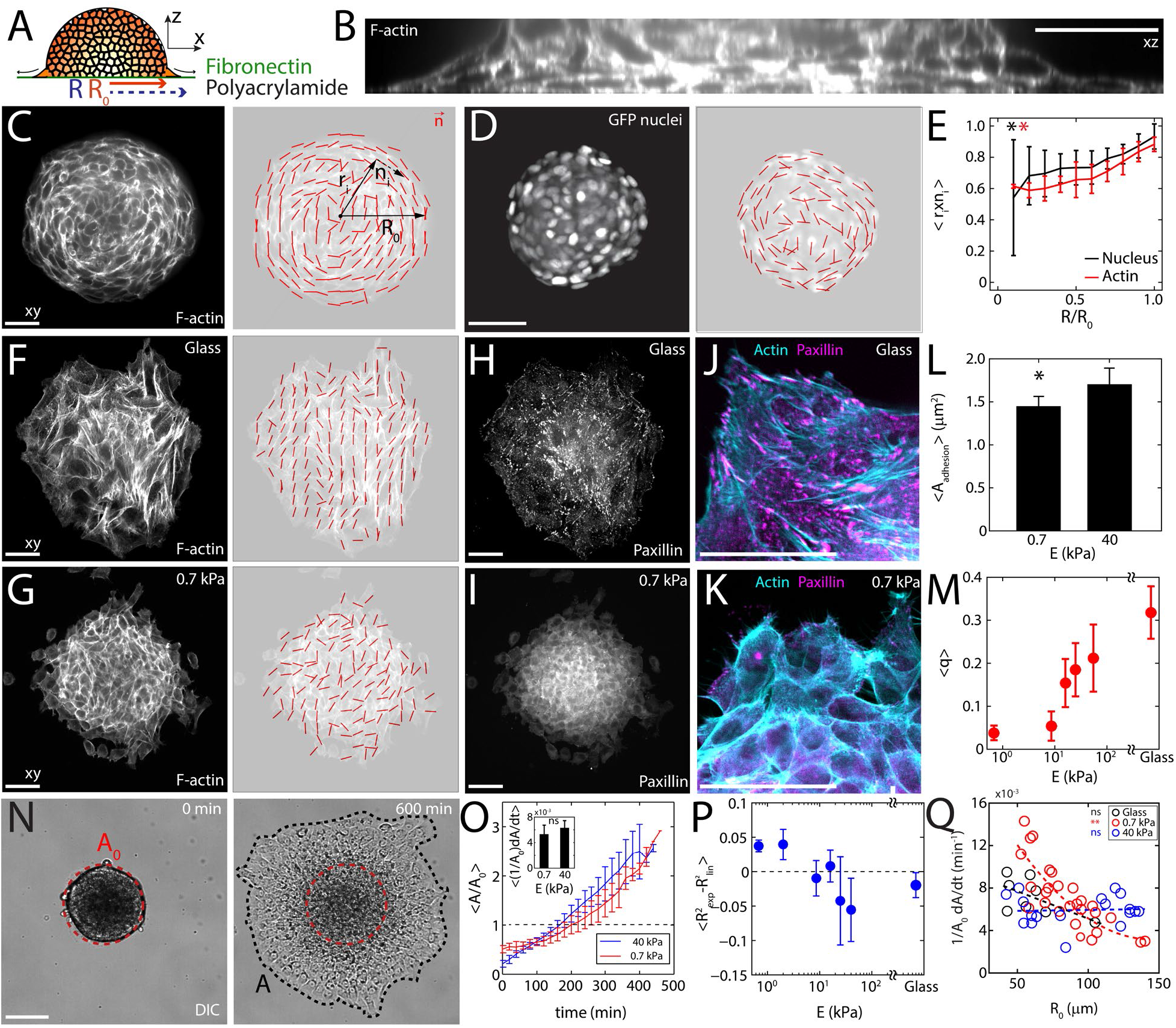
Substrate stiffness and aggregate size determine wetting dynamics. (A) Diagram of an aggregate spreading on a fibronectin-coated polyacrylamide gel. (B) Z-profile of F-actin stained aggregate adhered to glass. Fluorescently labelled F-actin within (C) an equatorial plane (z=10 μm) of an aggregate, red lines on an overlay indicate alignment. 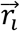 and 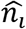 denote the distance of a nuclei from the center of an aggregate and its unit orientation vector respectively. (D) Organization of nuclei and corresponding alignment. (E) Mean orthoradial ordering in actin and nuclei as a function of distance from the aggregate center (n = 7). Fluorescently labeled F-actin within a monolayer spreading on glass (F), and 0.7 kPa gel (G) along with the corresponding alignment maps. Fluorescent Paxillin within the same monolayer on glass (H), and gel (I). Images showing focal adhesions in magenta and F-Actin in cyan for glass (J) and gel (K). (L) Average focal adhesion area (A_adhesion_) on 0.7 kPa and 40 kPa gels (n = 15). (M) Actin alignment parameter q = 〈*cos*^2^*θ*〉, as a function of substrate stiffness (n = 7, methods). (N) DIC image of an aggregate spreading on glass. A_0_ is the projected area of an undeformed aggregate, and A is the instantaneous area over which the monolayer has spread. (O) Normalized spread area (A/A_0_) as a function of time and stiffness. Inset displays the corresponding spreading rate 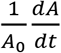 measured between A = A_0_ & A = 2A_0_ (n = 13 for 0.7 kPa and n = 10 for 40 kPa). (P) Difference in fitting R^2^ values as a function of stiffness. (Q) Spreading rate as a function of aggregate size for 0.7 kPa substrate (red, n = 27), 40 kPa substrate (blue, n = 18), and glass (black, n = 8). Scale bars are 50μm. *P < 0.05, **P < 0.01, ***P < 0.001. ns is non-significant. Error bars are mean +/− standard deviation.

## Traction stresses drive wetting on hard, but not soft substrates

Next, we measure the velocity field and stress field that characterizes the pattern of monolayer motion and force generation for a wide range of substrate stiffnesses. Particle Image Velocimetry (PIV) is used on basal (z=0 μm) images of cellular F-actin to measure the cumulative cell displacements which are normalized by the elapsed time yielding a velocity vector field, 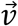 (Fig 2A). From this measurement, we observe that the velocity is principally oriented radially outwards, and the magnitude increases from the center of the aggregate to the periphery of the monolayer. Furthermore, like the strain sate, the gradient in velocity 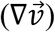 is invariant with the stiffness of the substrate (Fig 2D, 1O inset). Traction Force Microscopy (TFM) is used to measure the surface stresses 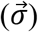 generated by cell motion (Fig 2B). In this case however, the stress field has a strong dependence on substrate stiffness. On rigid substrates (E > 8.6 kPa), stresses are concentrated at the periphery of the monolayer and are absent in the central region central to the aggregate (Fig 2C-F). By contrast, on compliant substrates (E < 2.8 kPa), stresses at the monolayer periphery are nominal, but elevated central to the aggregate. For intermediate substrate stiffness (E = 2.8 – 8.6 kPa), stress is uniformly distributed in space. Thus, while the velocity gradient is constant, the stress gradient varies continuously with substrate stiffness (Fig 2F). However, in all cases the stresses are oriented inwards, opposite to the direction of motion. Thus, the driving force for rapid wetting absent of elevated boundary stresses is unknown, and a localization of stresses away from boundaries indicate a fundamentally distinct mode of migration.

**Figure 2:**
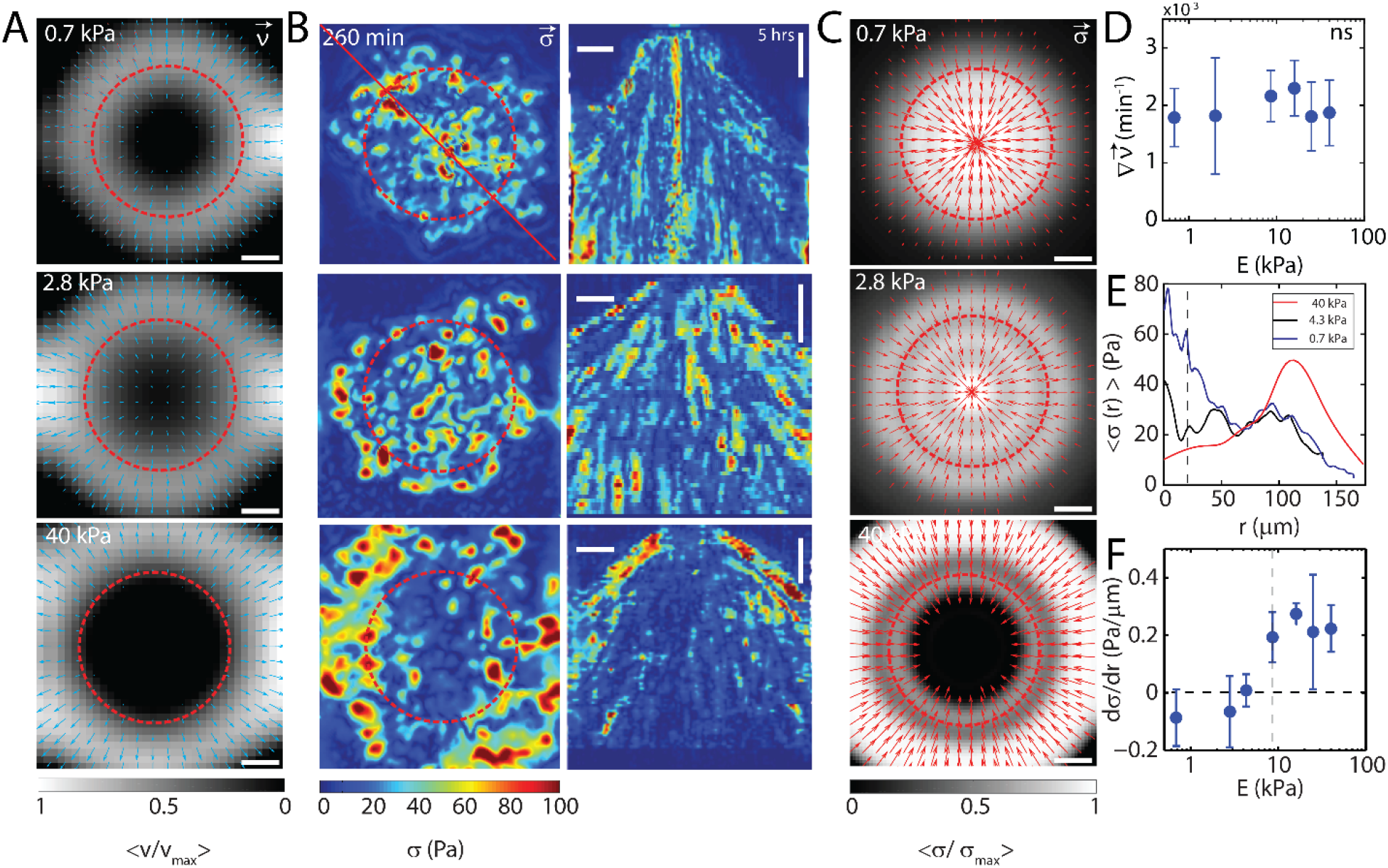
Traction stresses drive wetting on rigid but not compliant substrates. (A) Velocity fields 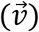 for cumulative displacement of cell motion. F-actin images are used for flow calculation. (B, left) Stresses (*σ*) calculated via TFM of aggregate spreading on substrates of stiffnesses 0.7 kPa (top), 2.8 kPa (middle), and 40 kPa (bottom). (B, right) Corresponding kymograph of data in (B, left). (C) Radial stresses 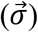 for the substrate stiffnesses in A. (D) Velocity gradient 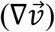 as calculated in (A), as a function of substrate stiffness, E (n = 12). (E) Radial stress distribution of a spread aggregate on substrates with stiffnesses 0.7kPa, 2.8kPa, and 40kPa (n = 1). (F) Stress gradient 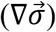 as a function of substrate stiffness, E (n = 12). ns is non-significant. Scale bar is 50 μm. Error bars are mean +/− standard deviation.

## Aggregate adhesion induces Laplace-like substrate indentation

The localization of stresses under the aggregate resembles the accumulation of pressure on a soft surface under an indenter, as described by contact mechanics. Indeed, the deformation is not limited to the plane, as the aggregate acts like a soft spherical punch deforming the substrate in the 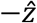 dimension (Fig 3A). We quantify the magnitude of the in-plane x-y deformation (dr) through PIV. For early times (*A* < *A*_0_), the contact area between the aggregate and the substrate increases, which is accompanied by a net inwards deformation of the substrate ∑ *dr* (Fig 3B). This can be compared to an increase in the contact area between a surface and a punch. At a contact angle of 90 degrees (*A* ≈ *A*_0_), dr is a maximum at the contact line and decreases towards the center (Fig 3C). By contrast, the out of plane indentation (z) is observed and maximized (z_max_) at the center of the aggregate (~0.5 μm), and decreases towards the contact line (Fig 3D). Thus, the magnitudes of x-y and z displacements are roughly equal across the surface of the aggregate. In addition, at the contact line, there is a positive deformation (in the 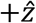 direction) of the gel, with the formation of a meniscus, suggesting strong adhesion (Fig 3D).

**Figure 3:**
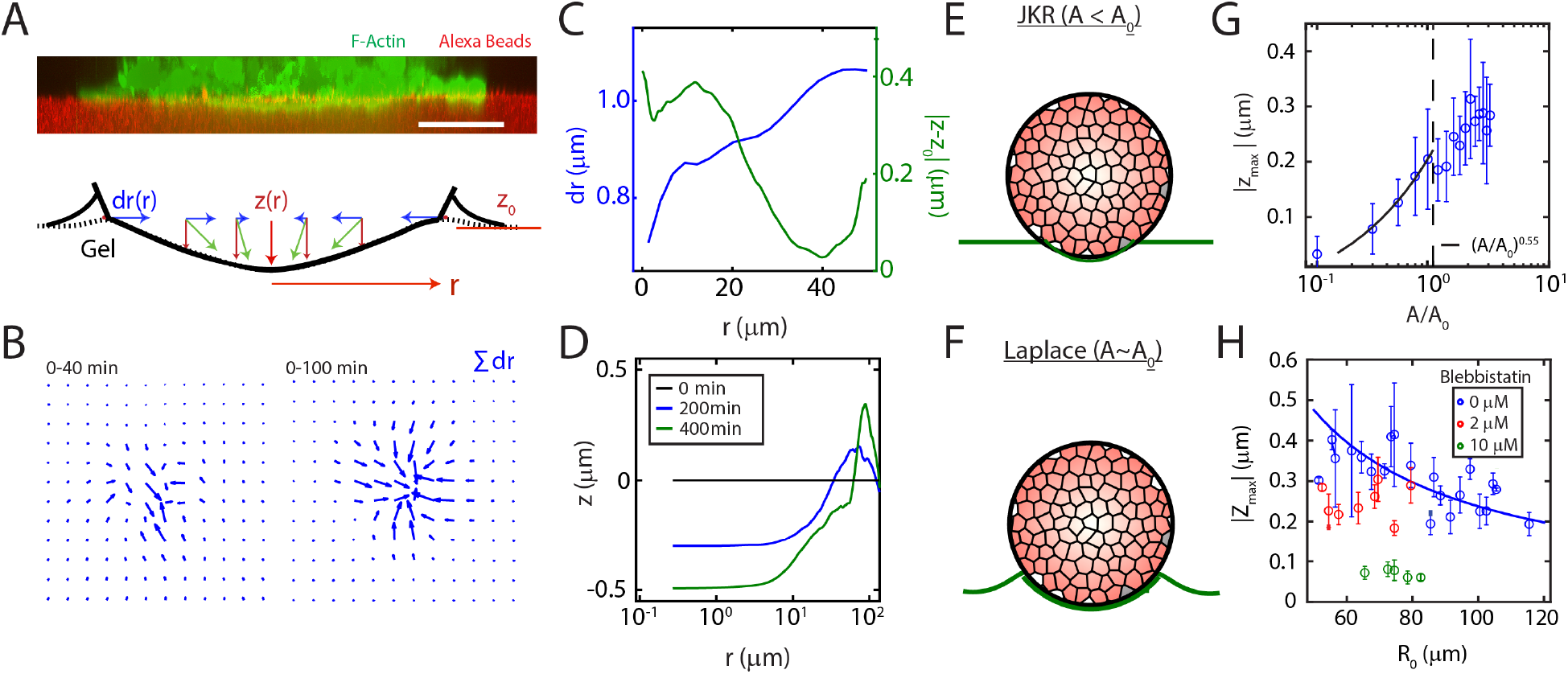
Adhesion and active stress induces elastic-like and fluid-like substrate indentations. (A, top) A reconstructed z-slice showing F-actin in a cell aggregate (green) indenting a 0.7 kPa substrate (red). (A, bottom) A schematic showing relative magnitudes of in plane (dr) and out of plane deformations (z). (B) Cumulative displacement field (∑ *dr*) over time during initial spreading (for A<A_0_) shows symmetric inward contractions of the substrate measured by PIV. (C) Magnitude of radial deformation dr, and z-indentation |z-z_0_|, at a single time (A~A_0_, approximately 90° contact angle) as a function of radial distance from the center of the aggregate r. (D) Z-indentation as a function of radial distance from the center of an aggregate. (E, G) Schematic of indentation of aggregate at early stage and absolute magnitude of indentation (z_max_) as a function of spread area (A< A_0_) (n = 7). (F, H) Schematic and maximum z-indentation (z_max_) as a function of aggregate size. Pharmacological treatment with Blebbistatin vanishes Laplace-like behavior. 0μM (n=75), 2μM (n=30), 10μM (n=20). Scale bar is 50 μm. Z_0_ is set to 0 for D, G, and H.

At early times (*A* < *A*_0_), indentation increases rapidly, reaching approximately 75% of the maximum by *A* ≈ *A*_0_. The Johnson Kendall Roberts (JKR) model in contact mechanics predicts the deformation of a surface caused by an elastic punch in the presence of adhesion^35^. In the original JKR model, the energy stored in deformation is greater than the adhesive energy, and a scaling of z ~ A is expected. However, we observe a roughly z ~ A^1/2^ relationship, consistent with a modified form of the JKR model, where the elasto-capillary length scale is comparable to the size of the punch, in which deformation and adhesive energies are comparable ^36^ (Figs 3E, G). This suggests that the early stages of aggregate spreading are driven by adhesion and an elastic solid interaction between the aggregate and the substrate.

At times *when A* ≈ *A*_0_ (contact angle ≈ 90°), the gel indentation has reached a maximum, |z_max_|. The law of Laplace relates the surface tension in a liquid with its internal pressure. It predicts that a droplet adhered to a substrate should induce an indentation that decays inversely as the size of the droplet. Indeed, dependent upon myosin activity, we find a |*z_max_*|~ 1/*R*_0_ relationship between the size of the aggregate and the indentation in the gel as has also been observed in passive droplets^36, 37, 38^ (Figs 3F, H, Supplementary Fig 5). This relationship is retained for a range of substrate stiffnesses, although the magnitude decreases with increasing substrate stiffness (Supplementary Fig 6). This suggests that the later stages of aggregate spreading are characterized by interactions between a fluid aggregate and an adhesive substrate. Equal in-plane and z-strains, as well as strong capillary interactions and a Laplace-like behavior are suggestive of strong surface tensions and internal pressures. Therefore, we next seek to measure aggregate surface tension directly and relate it to the size of the aggregate.

## Active stress yields size-dependent aggregate surface tension

Using micropipette aspiration we apply creep and stress relaxation tests on non-adherent aggregates to estimate the surface tension and other mechanical properties^29^. Briefly, a step in negative pressure is applied to the pipette, which draws the aggregate into the pipette a length, L (Fig 4A, Supplementary Figs 7,8, Supplementary Video 3). L will increase over time until the pressure is released, at which point L decreases (Fig 4B). By analyzing the flow of the aggregate using PIV we observe that the material flow is concentrated within ~50 μm of the pipette (Fig 4C, left). From the change of L in time, we estimate the viscosity (*η*) and the surface tension (*γ*) of the aggregate (Fig 4D,E, Supplementary Note 2). We find that *η* is independent of size and independent of applied force (Supplementary Fig 7). By contrast, *γ* increases with applied load (Fig 4F) and decreases with size (Fig 4G), consistent with the size-dependent indentation during aggregate adhesion. We confirm through finite-element modeling (FEM) that the observed change in surface tension does not correspond to a change in strain for aggregates of different sizes (Figs 4C right, 4G, Supplementary Figs 8, 9). This further rules-out a passive origin of size-dependent surface tension as the only way to replicate our experimental results; in FEM size-dependent surface stress must be incorporated explicitly. Furthermore, the magnitude of the *η* and *γ* are myosin-dependent, as both are reduced upon treatment with 50μM Blebbistatin, which inhibits myosin ATPase activity^39^ (Fig 4D,E,G, Supplementary Fig 8, Supplementary Video 4), also consistent with decreased indentation. In this case, *γ* is also no longer size-dependent, consistent with that of passive liquid droplets or DAH.

**Figure 4.**
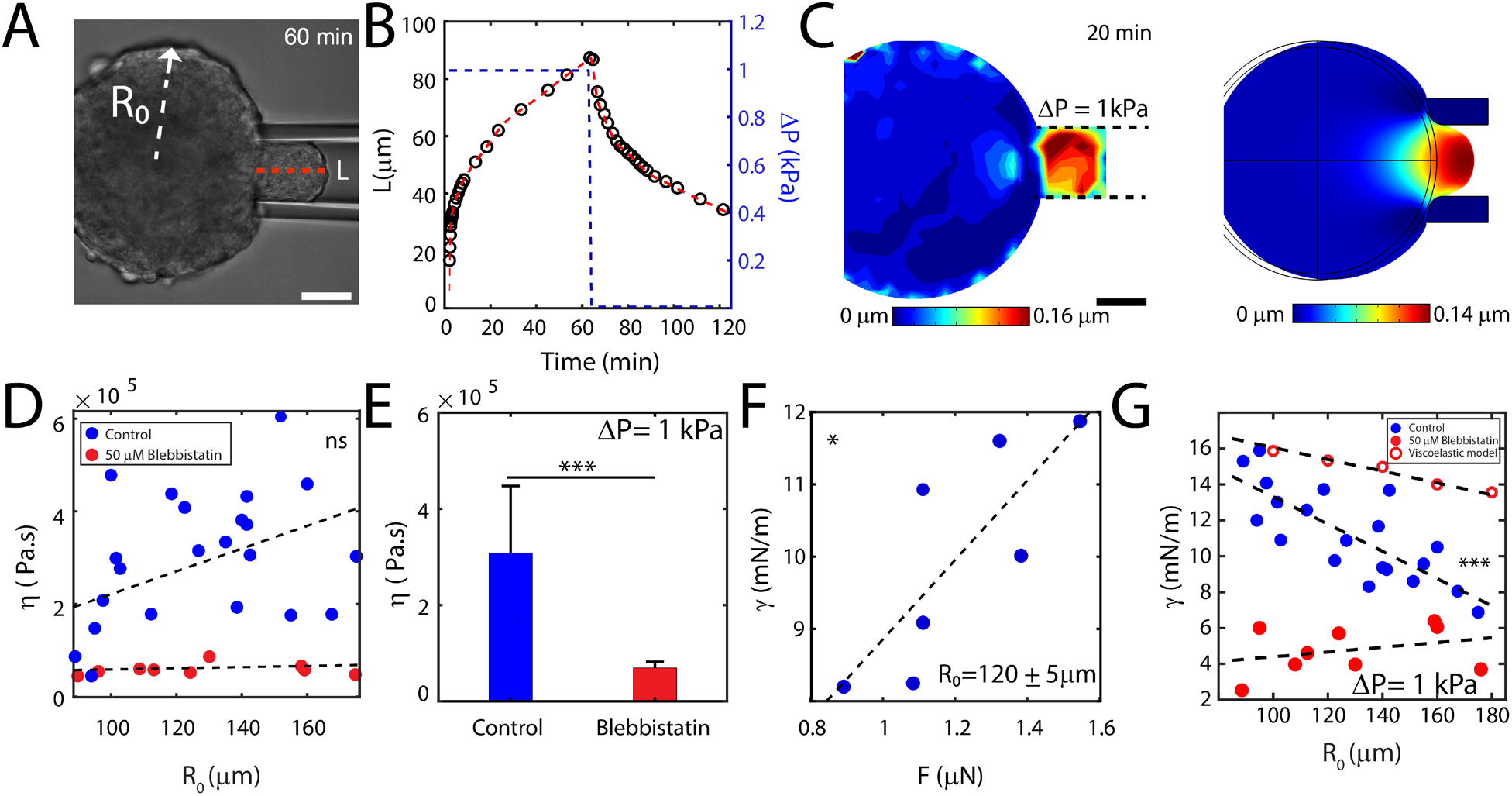
Aggregate surface tension is load and size-dependent. (A) DIC image of micropipette aspiration which shows an aggregate of radius R_0_ entering the pipette by a length L (indicated by dotted red line). (B) L over time (black dots, dotted red line). The pressure is quickly increased and then released at the apex of L (blue dotted line). (C) Colormap of displacement of cells in experiment (left) and in a finite element simulation of Burgers viscoelastic solid (right). (D) Viscosity (*η*) as a function of aggregate size, R_0_ (E) Viscosity in the presence (50 μM) and absence of Blebbistatin (n= 10 aggregates). (F) Surface tension (*γ*) as a function of applied pressure force (F), (n = 7) for aggregates of size R_0_=120μm ± 5μm. (G) Surface tension as a function of aggregate size (solid blue points), R_0_ (n=21) at a fixed applied pressure (ΔP=1kPa). Surface tension for aggregates in 50μM Blebbistatin (solid red points) (n = 8). Each data point is measured from one aggregate. Hollow red points show results of simulation of a Burgers viscoelastic solid. *P < 0.05, ***P < 0.001. ns is non-significant. Scale bar is 50 μm. Error bars are mean +/− standard deviation.

## Substrate stiffness differentiates compressible from incompressible cellular flows

As aggregate surface tension is sufficient to deform and indent soft substrates, and is size-dependent, we hypothesize that surface tension may explain the initial observation of a size-dependent spreading rate. To test this hypothesis, we seek to quantify the extent that pressure (elevated from myosin-dependent surface tension) impacts wetting. As pressure is difficult to measure during wetting, we first measure pressure in the non-adherent aggregate and relate it to cell density. Then, we measure changes in cell density during wetting, to infer the role of pressure as a motive force.

First, using the measured surface tension from micropipette aspiration, and the law of Laplace (P~γ/r), we calculate the internal pressure of the aggregates (Supplementary Fig 10). Separately, we find that the concentration of active myosin at the aggregate surface scales as ~1/*R*_0_ (Supplementary Fig 11). Further, we assume that γ is proportional to myosin concentration^40^. Therefore, substituting for γ in the law of Laplace, we find *P*~*V*^−2/_3_^, which agrees with experimental data (Supplementary Fig 10). By counting nuclei transfected with Nuclear-GFP, we estimate the population of cells (*N*) and cell density (*ρ* = *N/V*) within an aggregate (Supplementary Notes 4, 5). We find that 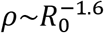 (Supplementary Fig 10). Substituting volume with density, we find a linear relationship between pressure and density (Supplementary Fig 10).

Second, we ask whether differences in density (and thereby pressure) may explain the difference in wetting dynamics based on substrate stiffness as initially observed (Fig 1O, Q). To do so, we use the principle of continuity and the balance of the influx current (I_1_) and the efflux current (I_2_) of cells (# *cells/min*) with areal cell density at the aggregate-substrate interface (*ρ_s_*, Fig 5A). We find that *ρ_s_* increases on soft gels, indicating I_1_ > I_2_ (Fig 5B, C). By contrast, *ρ_s_* does not increase significantly on glass (no clear relationship is observed on hard gels), indicating I_1_ ≤ I_2_. Furthermore, the efflux of cells (J_2_ = I_2_/A_0_) from the contact line, which is related to the spreading rate, is size-dependent (Fig 5D, Supplementary Notes 4, 6, Supplementary Figs 12). Both changes in density and size-dependent spreading rates may indicate a change in pressure. Thus, our hypothesis is strengthened, that on soft substrates (and for small aggregates), internal pressure may be involved in driving outward motion.

**Figure 5.**
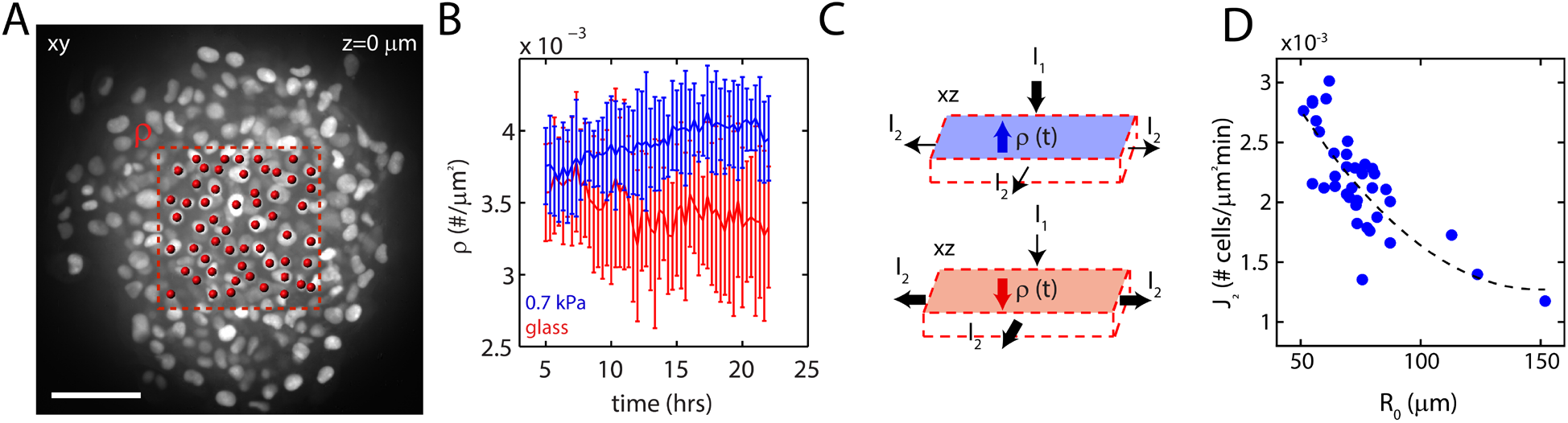
Substrate stiffness differentiates compressible from incompressible flows. (A) Schematic of a gaussian pillbox (red dotted line) to estimate the flux of cells entering from an aggregate towards the substrate and used to calculate surface number density of nuclei (ρ_s_). Red dots indicate spot-tracked nuclei. (B) Surface number density of cells (*ρ_s_*) as a function of time on glass and 0.7kPa substrates. (n = 10 each). (C) Schematic of the current of cells into the pillbox (I_1_) balanced with the efflux current (I_2_) that determines whether the density *ρ_s_* increases or decreases. (D) Cell efflux (J_2_) as a function of aggregate size (R_0_) on soft substrates (E=0.7kPa) (n = 38). Scale bar is 50μm.

## Density-dependent pressure is sufficient to drive size-dependent cellular flows

Thus far, we have inferred that pressure (elevated by surface tension) drives motion. To test the existence of pressure-driven flows more directly, we use a high energy laser to ablate cells at the surface of the non-adhered aggregate, in order to decrease aggregate surface tension and thereby alter pressure (Fig 6A, Methods). Immediately upon ablation, the surface of the aggregate retracts, indicating a drop in surface tension and the formation of an orthoradial velocity gradient at the surface (Fig 6B, Supplementary Video 5). Rearward surface flows are concomitant to forward flows of the core (center), suggesting the equilibration of internal stresses in response to the change in surface tension (Supplementary Fig 13). Furthermore, within 60s, the aggregate surface contracts to its original configuration, reversing flow of the core. Both forward and reverse behaviors are dependent upon myosin activity (Supplementary Fig 14). Thus, given the rapid response and large-scale flows, motion is likely pressure-driven.

**Figure 6:**
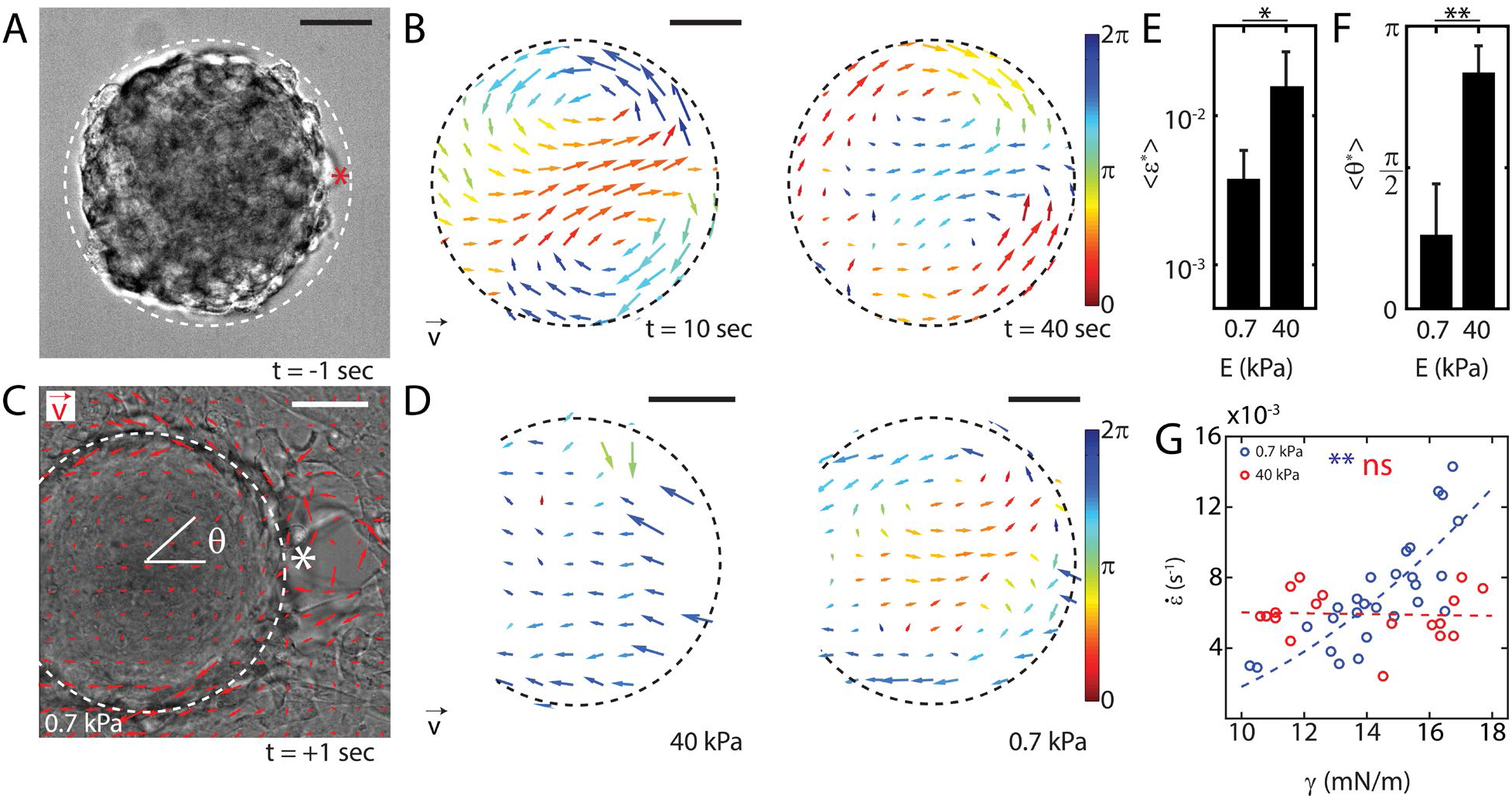
Perturbations in surface tension couple to pressure-driven cellular flows. (A) DIC image of a non-adherent aggregate immediately prior to ablation at the aggregate surface (t=0s). Location of the ablation is indicated by *. (B) Velocity field 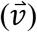 on surface and internal flows in a non-adhered aggregate, where displacement is accumulated over 10 seconds (n=5). Vector fields are color-coded by direction. (C) Aggregate adhered to 0.7 kPa gel, with displacement given by red PIV arrows, and the angle *θ*, with respect to the x-axis and ablation location. (D) PIV of an ablated aggregate to measure its response to step stress. (E) Strain (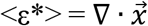, where 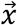 and 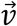 are related by a timescale) (n=8 each), and (F) direction of motion (<*θ*>) via PIV (n = 27). (G) Spreading rate 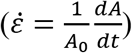 as a function of aggregate surface tension, *γ* on soft (blue) and hard (red) substrate. *P<0.05, **P<0.01. Scale bar is 50μm. Error bars are mean +/− standard deviation.

To understand if pressure-driven flows can be induced during migration, we ablate at the contact line of a spread aggregate – separating the interface between the aggregate and the monolayer (Fig 6C). In this case, we observe a retraction of both the aggregate and the monolayer, as measured by the resultant strain, as measured through PIV 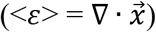 and the angle of displacement with respect to the location of ablation (<*θ*>), which depends upon substrate stiffness (Fig 6D). The monolayer retracts strongly on rigid substrates and only nominally on soft substrates. Retraction of the aggregate however, has two components: the displacement of the surface and the displacement of the core. The surface of the aggregate retracts for both soft and stiff substrates, indicating the presence of surface tension in both cases (Fig 6E). However, on rigid substrates, the core also retracts away from the ablated spot, while on soft substrates the core is driven towards the ablated spot (Fig 6F, Supplementary Fig 13, Supplementary Video 5). This may suggest an enhanced role for pressure on soft substrates in comparison to rigid substrates. Also, cleavage of spread aggregate adhesions (integrins & junctions) using trypsin shows strong retractions of the monolayer toward the aggregate on a 40 kPa substrate. However, on a 0.7 kPa substrate no retraction is observed (Supplementary Fig 15). This again, supports our hypothesis that the internal pressure of an aggregate may guide motion on soft substrates.

## Substrate stiffness differentiates pressure-driven from traction-driven flows

Having established experimental evidence that substrate stiffness may differentiate traction-based motion from pressure-based (and size-based) motion, we next show consistency of these results with models. First, we seek to relate the general dynamics of spreading on substrates of different stiffnesses to surface tension, through elasto-capillarity. Second, we explain the specific relationship between pressure and traction as compensatory mechanisms for driving robust wetting.

Balancing the gain in adhesion energy with dissipation through slipping explains the spreading on rigid substrates^34^, but is unable to capture the spreading behaviors observed on soft substrates. To explain the size-dependent spreading rate, we apply a model where additional dissipative effects occur through deformation of the substrate with a role for the active surface tension.

To capture the effect of the elastic response of the substrate, we explicitly account for the energy lost per unit time in deforming the substrate along with slipping losses and equate them to the energy gained per unit time due to spreading (Supplementary Note 1). Given that the energy changes can come from adhesion (E_adhesion_), viscous drag (E_viscous_), and deformation of the substrate (E_deformation_), we write

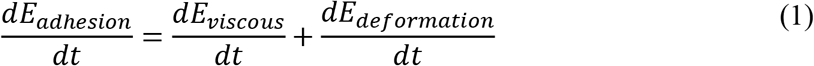

which leads to

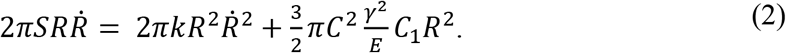

Here, S is the surface energy gained per unit area, R is the radius of the monolayer, k is the friction constant, γ is the surface tension of a freely suspended aggregate, E is the stiffness of the substrate, and C and C_1_ are constants. When the substrate stiffness is high, or at long time scales (A>A_0_) when there is a fully developed monolayer, the energy stored in the deformation of the substrate is negligible compared to its viscous counterpart. Neglecting the 2^nd^ term in the right-hand side of Equation 2 then gives a linear spreading behavior given by *R*^2^ = *Dt*, where D = S/k. This diffusive spreading behavior is consistent with the absence of a permeation width (Supplementary Fig 3) ^32^. At short time scales on soft substrates (A<A_0_), there is no monolayer. The energy gained by adhesion is lost in deforming the substrate. In this case we ignore viscous dissipation, leading to

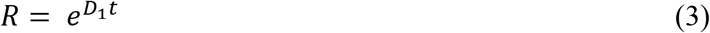

where D_1_ is 3πC^2^C_1_γ^2^/2E.

On soft substrates, the elasto-capillary length is comparable to the dimensions of the aggregate^38^, leading to an exponential spreading of the spread area. As substrate stiffness increases, the elasto-capillary length becomes vanishingly small and most of the dissipation occurs through slipping of the monolayer, causing the spreading rate to become linear as shown previously (Supplementary Fig 4)^34^. At long time scales, the dissipation through the monolayer is the leading dissipation term, again giving rise to a linear spreading rate.

Using the surface tension relationship from micropipette measurements, we plot the normalized spreading rate 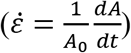 of an aggregate as a function of its surface tension (Fig 6G, Supplementary Fig 16). This is replotting of 1Q in terms of surface tension. As these measurements were performed on substrates of stiffness 0.7 kPa, the spreading is governed by Equation 3. We also note that the integrated forces applied by the substrate in this regime are comparable to the forces applied during micropipette aspiration (Fig 4, Supplementary Fig 17). Normalizing Equation 3 with the initial projected area of the aggregate, we estimate the time when the spreading monolayer had any given size. By comparing the times corresponding to A = A_0_ and A = 2A_0_, we find:

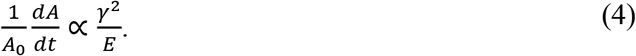

This indicates that the spreading rate increases quadratically with γ (Figure 6G). Replacing γ by *r_0_* we get a quadratic dependence of spreading rate on the initial radius of the aggregate (Supplementary Note 1). It has been shown in the case of stiff substrates that D increases linearly with the size of the aggregate, which implies that the spreading rate is independent of size. These behaviors on soft and stiff substrates are captured by fits in Fig. 1Q and 6G ^32^.

To determine how the balance between traction and pressure drives collective cell motion, we develop a vertex-based computational model to describe the spreading of the monolayer as a function of substrate stiffness ^41 42^. In this model, the monolayer is represented by a network of vertices, connected by tensed junctions that represent cell-cell interfaces ^43 44^. Each cell has a self-propelled force and exists in mechanical force balance with its neighbors. The influx of cells from the 3D aggregate (I_1_) is represented by cell addition localized to a fixed, central and circular 2D region interior to the boundary (Supplementary Note 7). The cell incorporation rate within this region is independent of substrate stiffness (E, unitless) although the single cell self-propulsion force and cell-substrate friction coefficient correlates positively with E (Supplementary Notes 7, 8, Supplementary Figs 18, 19, Table 4). With these model properties, the monolayer spreads and traction stresses grow to localize either to the monolayer boundary (E > 15) or to the interior (E < 10) consistent with our experiments (Fig 7A-B, Supplementary Fig 18). Respectively, internal pressures either decay (E >15) or grow (E < 15) in time (Fig 7C, D) positively correlated to the density of cells within the monolayer (Supplementary Fig 18). Thus, motion may be principally traction-driven (E >15) or guided by pressure (E <15) where traction stresses are nominal (Fig 7E-G).

**Figure 7:**
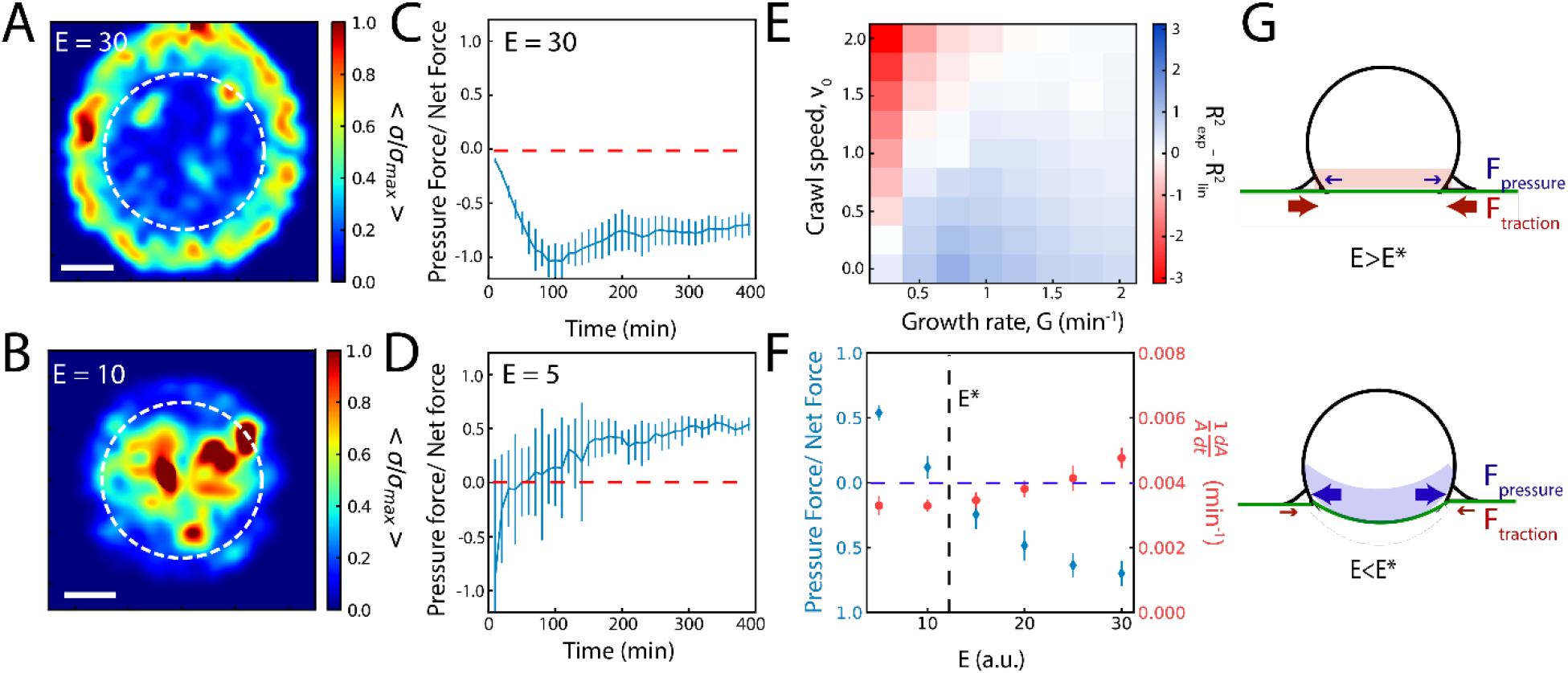
Pressure compensates for traction as an outward motive force. Traction force results from Vertex Model for soft, (A) E = 10 and stiff, (B) E = 30 substrates. Area outlined by dashed white line indicates the region of cell addition. Scale bar is 20 μm. (C) On rigid substrates (E = 30), friction is high, and self-propulsion accounts for most of the total traction force that drives motion. (D) On soft substrates (E = 5) friction is low, and pressure accounts for most of the total traction force that drives motion. (E) The balance of crawling and growth (cell addition, representing I_1_) reproduces exponential and linear behaviors. (F) The balance of pressure to self-propelled forces changes as a function of substrate rigidity (E), with nominal changes to spreading rate 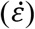. (G) Schematic for pressure-driven and traction-driven flows, which depends upon the stiffness of the substrate, E. Error bars are mean +/− standard deviation.

## Discussion

In this work, we use a combination of experimental force measurements, computational models, and theory to understand the how the balance between interfacial tensions out of equilibrium and substrate elasticity drives the wetting of cell aggregates.

In cell aggregates, the non-equilibrium activity of molecular motors endows the surface tension with a dependence upon aggregate size. As a result of this dependence, the balance between tensions and substrate elasticity is not constant and determines the motive forces that drive the dynamics of wetting.

Upon wetting of rigid substrates, mature focal adhesions and organized F-actin generate large traction stresses at the boundary of the monolayer that drives outward expansion. In this case, the boundary pulls the monolayer outward, and the internal density of the aggregate remains nearly constant or decreases. By contrast, upon wetting a compliant substrate, boundary stresses are attenuated due to the inability of immature focal adhesions and a disordered F-actin cytoskeleton to generate significant traction. Yet, for small aggregates in particular, the monolayer expands at a comparable rate. In this case, the accumulation of cells upon the substrate increases local cell density and cell density determines the internal pressure. With low friction, this pressure in turn can guide outward cellular flows (Supplementary Video 6) aiding in the expansion of the monolayer film (“traction-independent flow”). By extension, as surface tension depends upon aggregate size, so does the motion of cells from the aggregate. Thus, the wetting of aggregates utilizes distinct driving mechanisms to adapt to the mechanics of their surroundings.

In the present study, the presumption of a uniform pressure is sufficient to reproduce experimental results. However, while not modeled explicitly, the spatial variation of actin alignment within the aggregate (Fig 1) is consistent with a gradient in internal pressure^45^ which may further contribute to outward flows.

These results indicate that surface-tension driven pressure and traction stresses are compensatory mechanisms during the spreading of cell aggregates. Further, these results suggest that bulk properties of tissues can influence the collective motion of cells, providing new insights for the study of early development and cancer metastasis.

## Acknowledgements

We acknowledge funding ARO MURI W911NF-14-1-0403 to MM & APT as well as NIH RO1 GM126256 to MM and NIH U54 CA209992 to MM & MSY. MFS is supported by a UK Engineering and Physical Sciences Research Council (EPSRC) PhD studentship at University College London (UCL). SB acknowledges support from Royal Society grant # URF\R1\180187. SB and MPM also acknowledge support from Human Frontiers Science Program (HFSP) grant # RGY0073/2018. Any opinion, findings, and conclusions or recommendations expressed in this material are those of the authors(s) and do not necessarily reflect the views of the NSF, NIH, HFSP, Royal Society, or EPSRC. We would like to thank Prof. Erdem Karatekin for help with micropipette measurements. We would also like to thank Dr. Karine Guevorkian, and Prof. Francoise Brochard-Wyart for constructive discussions.

## Author Contributions

MPM designed and conceived the experimental work. SB designed and conceived the computational model. VY conceived the theoretical model. MSY, SA, GJ, SA, and YE acquired experimental data. MPM, VY, SA, and MSY analyzed experimental data. MFS designed and implemented the active vertex model and performed simulations. SA implemented and performed finite element simulations. MPM and VY wrote the paper, with help from MSY, APT, SA, MFS and SB.

## Competing Financial Interests

The authors declare no competing financial interests.

## Materials

### Cell Culture and Aggregates Preparation

We use murine sarcoma E-cadherin expressing cells with a doubling time of approximately 24 hours^1^. Cells are cultured at 37 °C under 95% air/ 5% CO_2_ atmosphere in culture medium consisting of Dulbecco’s Modified Eagle Medium (DMEM) enriched with 10% calf serum and 5% penicillin streptomycin. Cell aggregates are prepared from confluent cell cultures using a suspension-spinning method. Aggregates ranging from 80 to 450 μm in diameter (Supplementary Fig 2) are obtained from 5 mL of cell suspension in CO_2_-equilibrated culture medium at a concentration of 4×10^5^ cells per mL in 25 mL Erlenmeyer flasks, which are placed on a gyratory orbital shaker at 75 rpm at 37 °C for 30-50 hrs. The flasks are pretreated with 2% dimethylchlorosilane in chloroform to prevent adhesion of cells to the glass surface.

### Cell transfection for Actin

Murine sarcoma S-180 cells were stably transfected with plasmid construct encoding for F-TRActin-EGFP (Addgene^®^ Plasmid #58473). Briefly, cells were transfected using FuGENE^®^HD Transfection reagents where 2 μg of the DNA plasmid was added to the transfection reagent and added to a cell dish. Cells were incubated for over 24 hrs with the plasmid to complete the transfection process. Following 1 week of incubation with a selection media containing G418 (Mirus Bio LLC) at 1mg/mL, population of cells were isolated and flow sorted with BD FACSAria II flow cytometer. The isolated population were cultured and expanded in DMEM (supplemented with 10%FBS, 1%Pen/strep) and used for experiments.

### Cell transfection for Nuclear-GFP

GFP tagged with a NLS (nuclear localization sequence) was introduced in a lentivirus plasmid using Gibson assembly. GFP-NLS expression is driven in this plasmid by the PGK promoter. For virus production, envelope plasmid pMD2.G, packaging plasmid psPAX2, and library plasmid were added at ratios of 1:1:2.5, and then polyethyleneimine (PEI) was added and mixed well. The solution stood at room temperature for 5 min, and then the mixture was added into 80-90% confluent HEK293FT cells and mixed well by gently agitating the plates. Six hours post-transfection, fresh DMEM supplemented with 10% FBS and 1% Pen/Strep was added to replace the transfection media. Virus-containing supernatant was collected at 48 hours, and was centrifuged at 1500 g for 10 min to remove the cell debris; samples were aliquoted and stored at −80 °C. Lentivirus was added to the media of sarcoma cells (30% confluency) with polybrene (Millipore-sigma TR-1003-G) at a concentration of 5μg/ml. After 2 days of incubation GFP-positive cells were sorted on a BD FACSAria II.

### Immunofluorescence staining

Adherent and non-adherent aggregates were fixed with a solution of 4% paraformaldehyde in 1X PBS for 15 min and permeabilized with 0.2% Triton X-100 for 20 min. The cells were then blocked with 2% bovine serum albumin (BSA) in 1X PBS (PBS-2%BSA) at room temperature for 1 h, incubated with primary antibodies for Paxillin (Recombinant Anti-Paxillin antibody [Y113], Abcam ab32084; 1:250 dilution) and Myosin (Phospho-Myosin Light Chain 2 (Ser19) Antibody, Cell Signaling Technologies^®^ (#3671); 1:250 dilution) in PBS-2%BSA for 48 hrs at 4°C. The sample were incubated for 48 hrs at 4°C for secondary antibodies Alexa Fluor 647 (Donkey anti-Rabbit, Abcam ab 150075, 1:500 dilution) and Alexa Fluor 555 (Anti-mouse IgG (H+L), F(ab’)2 Fragment, 1:500 dilution). Phalloidin staining was performed with Alexa Fluor 488 Phalloidin (Life Technologies; 1:200) diluted in PBS with 2% BSA for 48 hrs at 4°C. Images were taken with 2X (air), 40X and 60X oil immersion objective. Analysis was performed with ImageJ and Imaris (Bitplane).

### Blebbistatin treatment

Myosin II inhibitor ((-) Blebbistatin) was purchased from Sigma Aldrich (B0560) and reconstituted in dimethyl sulfoxide (DMSO) at high stock concentration of 17.1mM. Different dilution concentrations were used for experiments as 10 μM, 25 μM and 50 μM unless otherwise noted.

### Polyacrylamide Gel Preparation

Traction force microscopy experiments were carried out on polyacrylamide (PA) substrates polymerized onto 25mm diameter (#1.5, Dow Corning) coverslips. Briefly, the coverslips are treated with a combination of aminopropylsilane (Sigma Aldrich) and glutaraldehyde (Electron Microscopy Sciences) to make the surface reactive to the acrylamide. The ratios of polyacrylamide to bis-acrylamide for the gels used in this study are 7.5%:0.03% (0.7kPa), 7.5%:0.1% (2.8kPa), 7.5%:0.153% (4.3kPa), 7.5%:0.3% (8.6kPa), 12%:0.145% (16kPa), 12%:0.19% (20kPa), 12%:0.25% (25kPa), 12%:0.46% (40kPa) and 12%:0.6% (55kPa). A concentration of 0.05% w/v ammonium persulfate (Fisher BioReagents) and 20nM beads (Molecular Probes) are embedded in the gel mixture prior to polymerization. A 15 μl volume of the polyacrylamide solution is added to the coverslip and covered with another coverslip, which has been made hydrophobic through treatment with Rain-X^®^. The gels are polymerized on the coverslips for 30 minutes at room temperature. The gels are then reacted with the standard 1mg/mL Sulfo-SANPAH (Thermo Fisher Scientific, 22589) ^46^. The surface of the gels is then coated with fibronectin (F002, Sigma-Aldrich) with 1 mg/ml concentration (Supplementary Figs 1, Table 1, 2). Lower concentrations of fibronectin changes dynamics of wetting to dewetting and partial wetting conditions. The reaction proceeds for 12 hrs overnight incubation in the dark, and the coverslips are then rinsed and stored in 1X PBS.

### Traction force microscopy

Traction force microscopy is used to measure the forces exerted by cells on the substrate. ‘Force-loaded’ images (with cells) of the beads embedded in the polyacrylamide gels were obtained using a 60× oil-immersion objective (Leica Microsystems). The ‘null-force’ image was obtained at the end of each experiment by adding trypsin to the cells for 1 h. If the wound was exceedingly large (that is, of the size of the image) the ablated image was used for the null-force image. Images were aligned to correct drift (StackReg for ImageJ) and compared to the reference image using PIV software (http://www.oceanwave.jp/softwares/mpiv/) in MATLAB to produce a grid spacing of 7.0 μm. Forces can be reliably measured between 4.3 and 40 kPa and are calculated using custom-written code by Ulrich Schwarz.

### Micropipette aspiration

We performed the aspiration of the aggregates using pipettes with diameters 3–5 times that of a single cell (40–70 um). The micropipettes were fabricated by pulling borosilicate glass pipettes, with outer and inner initial diameters of 1mm and 0.5mm respectively, with a laser-based puller and desired diameter was achieved by cutting with a quartz tile. To prevent adhesion of the cells to the micropipette walls, the pipettes were incubated in 0.1mg/mL PolyEthyleneGlycol-PolyLysine (PLL-PEG) in HEPES solution (pH 7.3) for 1 h. The observation chamber consisted of microscopic slides. Aggregates were then suspended in CO_2_ equilibrated culture medium the pipette was introduced into the chamber. To prevent evaporation, the open end was sealed with mineral oil. A range of pressures (P: 500 Pa to 1.5 kPa) was attained by vertically displacing a water reservoir, connected to the pipette, with respect to the observation chamber. Aspirated aggregates were visualized on an inverted microscope equipped with a 20X air objective. Movies of the advancement of the aggregates inside the pipette were recorded with a CCD camera with a 5–30 s interval. All experiments were performed in physiological temperatures of 37°C.

### Particle Tracking

Individual nuclei are monitored by spot tracking of the center of fluorescence intensity in Imaris (Bitplane). For quantifying alignment of nuclei in an aggregate, the edges of nuclei were marked manually. The orientation was then detected by fitting the enclosed region with an ellipse. The direction of orientation is then the principal axis of the fitted ellipse.

### Director field calculations

The director field 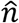 is calculated using custom MATLAB^®^ code as described previously.^47^ Briefly, a director field is created from images of fluorescently labeled F-actin and the images are divided into small, overlapping 8.6 μm by 8.6 μm windows, and the local orientation director is calculated for each window^48^. Each window is then Gaussian filtered and transformed into Fourier space using a 2D fast Fourier Transform (FFT). Then, the axis of the least second moment is calculated from the second order central moments of the transformed window. The angle of the local F-actin director is defined as orthogonal to this axis.

### Nematic order calculation

The local degree of alignment is calculated between adjacent windows within 3×3 kernels. The local nematic order is calculated for the central window in each kernel using the modified order parameter equation *q* = 〈*cos*^2^*θ*〉, where θ is the difference in F-actin orientation between the central window and the 8 surrounding windows. This process is repeated for all possible 3×3 kernels over an image, yielding a nematic director field with defined director magnitude and orientation for each window over an image. Perfect alignment between adjacent regions within an F-actin network results in an order parameter equal to one.

### Particle Image Velocimetry

Particle image velocimetry (PIV) is applied in MATLAB (MathWorks^®^) to fluorescent F-actin images, (mPIV, https://www.mn.uio.no/math/english/people/aca/jks/matpiv/) yielding displacement & velocity vector fields.

### Laser ablation

Laser ablation was performed using a 435 nm laser (Andor Technology). A 20X (air) objective (Leica Microsystems) was used for ablation and the laser power was held at 90%. The sample were ablated along a straight line of approximately 100μm length with 10 spots of equal spacing. Images were acquired before and after ablation for 1 s intervals for 16 min with a confocal microscope (ANDOR, Oxford Technologies, Belfast, Northern Ireland).

### Focal adhesion size calculation

Spatial dimension of focal adhesion spots was determined using point detection tool in Imaris software (Bitplane).

### Statistical tests

All statistical comparisons between two distributions were done with a two-sided *t*-test. When distributions are presented as a single value with error bars, the value is the mean of the distribution, and the error bars are the standard deviations. We use the symbols *, ** and *** for *p*<0.05, 0.01 and 0.001, respectively. When fitting lines to data, we quote the *p*-value as significance values to rejections of the null hypothesis.

### Cell-based computational model

For a full description of the computational active vertex model see the Supplementary Information (Supplementary Notes 7 & 8).

### Data availability

Data that support plots and other findings within this manuscript are available from the corresponding author upon reasonable request.

### Code availability

Custom codes that were used to analyze experimental data within this manuscript are available from the corresponding author upon reasonable request.

## Notes

### Competing Interest Statement

The authors have declared no competing interest.

